# An adaptable chromosome preparation methodology for use in invertebrate research organisms

**DOI:** 10.1101/226084

**Authors:** Longhua Guo, Alice Accorsi, Shuonan He, Carlos Guerrero-Hernández, Shamilene Sivagnanam, Sean McKinney, Matthew Gibson, Alejandro Sánchez Alvarado

## Abstract

**Background:** The ability to efficiently visualize and manipulate chromosomes is fundamental to understand the genetic architecture of organisms. This can be challenging for many invertebrates because conventional chromosome preparation protocols deal primarily hypotonic conditions tailored for use with vertebrates and rely extensively on cultured cells which are rarely available for invertebrates. Hence, a simple and inexpensive chromosome preparation protocol, adaptable to multiple invertebrate model species is needed.

**Results:** We optimized a chromosome preparation protocol and applied it to several planarian species (Phylum: Platyhelminthes), the freshwater apple snail *Pomacea canaliculata* (Phylum: Mollusca), and the starlet sea anemone *Nematostella vectensis* (Phylum: Cnidaria). We showed that both mitotically active adult tissues and embryos can be used as sources of metaphase chromosomes, expanding the potential use of this technique to invertebrates lacking cell lines and/or with limited access to the complete life cycle. Simple hypotonic treatment with DI water was sufficient for karyotyping. The karyotypes we obtained allowed the identification of differences in ploidy and chromosome architecture among otherwise morphologically indistinguishable organisms, as in the case of a mixed population of planarians collected in the wild. Furthermore, we showed that in all tested organisms representing three different phyla, this protocol can be effectively coupled with downstream applications, such as chromosome fluorescent *in situ* hybridization.

**Conclusion:** The simple and inexpensive chromosome preparation protocol reported here can be readily adapted to new invertebrate research organisms in order to expand and accelerate the discovery of new biology in understudied branches of the tree of life.

## Background

Chromosome preparation is instrumental to establish new research organisms and to study genetics, genomics and evolution. As genome sequencing has become more and more affordable, the biological research community has witnessed in the last decade alone an explosive number of understudied organisms being sequenced and studied. Identifying the basic chromosome composition of an organism sets a foundation to query its unique biological attributes [1, 2]. Behaviors of cells and chromosomes can be fundamentally different in animals that are stable diploids, stable polyploids or possessing mixed-ploidy. Hence, experimental observations at genetic/genomic, tissue, and organismal levels need to be interpreted accordingly.

Isolated chromosomes can be used directly to study multiple questions in genetics and epigenetics. For example, they can be used to collapse scaffolds into chromosomal level genome assembly [3], to detect insertion or deletion of large fragments of DNA [4], to visualize physical positions of multiple genes [5, 6], to localize histone modifications or to identify DNA regions interacting with chromatin regulators [7, 8]. Moreover, genomes assembled into chromosomes can be used for comparative mapping to study chromosome and genome evolution [9–12].

For communities interested in evolutionary genetics and evolutionary developmental biology, it is favorable to learn about chromosome compositions of multiple species of interest. Establishing a new research organism is fraught with many technical challenges and biological unknowns. The development of freshwater planarians as a popular invertebrate for studying regeneration and adult stem cells illustrates many of these challenges and opportunities [13–17]. Existing literature since early 20^th^ century on chromosome compositions of freshwater planarians showcased the dynamic changes of ploidy within a species and the wide range of chromosome numbers across closely related species [18]. While these phenomena provide opportunities to study genetics and evolution [19], it also demonstrated the necessity of a standardized chromosome preparation protocol to identify animal ploidy [14].

An optimized chromosomal preparation protocol that can be easily adapted to multiple species would be advantageous. To establish a new research organism, researchers often run into problems with tool-building because of basic biological differences that require significant adaptations to many “standard” protocols. Conventional karyotyping protocols in mammalian cells involve tissue disassociation and cell culture [20] which are not applicable to most invertebrate research organisms. Furthermore, osmolality, temperature, time and centrifugation speed are all complex factors to optimize for karyotyping an invertebrate species of interest. This kind of complexity and need of an optimized protocol is clearly demonstrated by the various hypotonic reagents, time and temperatures used in chromosome preparations in the planarian literature [21].

Here, we optimized a chromosome preparation protocol for the purpose of chromosome visualization and manipulation in freshwater planarians (Phylum: Platyhelminthes), and two other emerging research organisms, *Pomacea canaliculata* (Phylum: Mollusca) and *Nematostella vectensis* (Phylum: Cnidaria). We demonstrated that: 1) both adult and embryonic tissues, treated with colchicine, can provide an adequate number of mitotic cells, 2) DI water is a convenient hypotonic reagent; and 3) there is no need of tissue dissociation, centrifugation or cell culture. We also demonstrated that the chromosomes prepared with this protocol are suitable for downstream applications, such as fluorescent *in situ* hybridization (FISH) in all the tested species. Our protocol is simple, inexpensive and potentially can be adapted to a wide variety of invertebrate research organisms.

## Methods

### Planarians

Both sexual and asexual strains of *Schmidtea mediterranea*, in addition to asexual strains of *Dugesia japonica, Phagocata velata* and wild planarians collected in the field (Sardinia, Italy) were maintained in the dark at 21–22 °C in 1× Montjuïc water (1.6 mM NaCl, 1.0 mM CaCl_2_, 1.0 mM MgSO_4_, 0.1 mM MgCl_2_, 0.1 mM KCl, and 1.2 mM NaHCO_3_ in DI H_2_O, pH 6.9–8.1) [22]. All animals were fed organic beef liver paste twice per week, and the culture water was exchanged after each feeding. The animals were starved for 1 week before experiments.

#### Pomacea canaliculata

Specimens of the apple snail *P. canaliculata* were maintained in the lab at 26-27 °C with a 14:10 h light:dark cycle. The animals were housed in tanks filled with artificial freshwater (2.7 mM CaCl_2_, 0.8 mM MgSO_4_, 1.8 mM NaHCO_3_, 1:5000 Remineralize Balanced Minerals in Liquid Form [Brightwell Aquatics]). The water was changed twice per week, followed by cleansing of the tanks. The snails were fed a variety of lettuce leaves. Laid embryos were collected daily and stored in dry conditions.

### Nematostella vectensis

Colonies of the sea anemone *N. vectensis* were maintained at 18-20 °C in 1:3 dilution of artificial sea water (12 parts per thousand [ppt] of Instant Ocean Sea Salts) and were fed the larvae of the brine shrimp *Artemia salina* two to five times per week. Spawning induction of sexually mature individuals, egg de-jellying treatment, and fertilization were carried out as previously described [23].

### Planarian sample preparation for chromosome spreads

The tails of planarians were amputated using a blade, and the regenerating tail fragments were incubated in 0.25% colchicine (Sigma Aldrich) in 1× Montjuïc water for 0 or 6 h in the dark. To test the efficiency of colchicine treatment, some tail fragments were incubated for 0 or 6 h in 1× Montjuïc water without colchicine.

### *P. canaliculata* sample preparation for chromosome spreads

*P. canaliculata* embryos 5, 6, 8 or 10 days post-fertilization (dpf) were collected, and both the external egg shell and the bright pink perivitelline fluid were removed with forceps. The resulting embryos were incubated in 0.25% colchicine in Pc-embryonic salt solution (33.3 mM NaCl, 6 mM KCl, 6.7 mM CaCl2, 3.3 mM MgCl2, 1.67 mM HEPES,) for 6 or 24 h in the dark. To test the efficiency of colchicine treatment, some embryos were incubated in *Pc*-embryonic salt solution without colchicine.

Adult snails (2.8–3.2 cm in diameter) were incubated in 0.25% colchicine in artificial freshwater for 6, 24, 48 or 72 h. After the incubation, small fragments of the organs of interest were dissected with scissors.

### *N. vectensis* sample preparation for chromosome spreads

Fertilized *N. vectensis* embryos were cultured for approximately 5 h post-fertilization (hpf). At 64- or 128-cell stage, around 200 embryos were transferred to a sterile 50-mm Petri dish. They were rinsed twice with 12 ppt artificial seawater to remove extra sperm and incubated in freshly made 0.04% colchicine in 12 ppt artificial seawater for 45 min with gentle rotation in the dark.

### Preparation of siliconized coverslips

Siliconized coverslips (22 × 22 mm) were submerged in Sigmacote^®^ (Sigma Aldrich) for a few sec in a fume hood. The coverslips were then propped up vertically for 10 min to dry, rinsed in DI H_2_O for a few sec, and propped up vertically to dry.

### Chromosome spread preparation for planarians

Tail fragments were placed in a Petri dish after colchicine treatment and rinsed with DI H_2_O. The tissue was punctured using forceps or needles to increase its permeability and then incubated in DI H_2_O for 20 min at room temperature (RT). The samples were fixed with freshly made Carnoy’s fixative (3:1 dilution of methanol:acetic acid) for 30 min on ice. A small portion of the tail fragment was then placed onto a slide using a pair of forceps or a pipette. The sample was soaked in a drop of 60% acetic acid (10–20 μl) and incubated for 5 min. A siliconized coverslip was placed on top of the tissue. The tissue was then squashed by applying constant pressure to the coverslip for a few sec to create a single layer of nuclei. Care was taken to keep the coverslip from shifting laterally during the application of the coverslip. After an overnight (ON) incubation at 4 °C, the slides were chilled on dry ice for 10–20 min. While the slides were still sitting on the dry ice, coverslips were quickly removed using a blade. The blade was inserted between the slide and one corner of the coverslip and used as a lever for removing the coverslip, with care taken to not scratch the slide where the sample is sitting. The slides were subsequently returned to RT, rinsed 3 times with 1× phosphate-buffer saline (PBS), and stained with a 1:5000 dilution of 4’,6-diamidino-2-phenylindole (DAPI) in 1× PBS for 10 min at RT. The slides were then rinsed twice with 1× PBS for 5 min each and mounted with either Prolong Diamond (Molecular Probes) or Vectashield (Vector Laboratories) mounting media. The slides were stored at 4 °C.

### Chromosome spread preparation for *P. canaliculata*

All procedures were the same as those described in “Planarian Section” except for the following modifications:

- The adult tissues were punctured using a needle, but not the embryos.
- Before placing tissues on slides, the older embryos (10 dpf) were cut in half and only the anterior portion was squashed.

### Chromosome spread preparation for *N. vectensis*

All procedures were the same as those described in “Planarian Section” except for the following modifications:

- Because *Nematostella* embryos are extremely fragile during early cleavage stages, significant caution was taken when changing solutions. Specifically, solutions were added slowly dropwise and enough solution was left in the Petri dish to keep the embryos fully submerged to avoid blastomeres burst as a result of the surface tension exerted by the solutions.
- The embryos were not punctured with a needle and the initial DI H2_O_ incubation was performed for only 5 min due to the fragility of the blastomeres.
- After 5 min in 60% acetic acid, the weight of the siliconized coverslip was sufficient to flatten the cells, and no additional pressure was applied.

### Image acquisition and processing

The spreads were imaged on a Zeiss LSM 780 or Zeiss LSM 700 confocal microscope using 63× or 100× magnification lens. The acquired images were processed and karyograms were generated using Photoshop and FiJi.

### Protocol optimization tips

To adapt this protocol for use in other research organisms, it is important to consider the following key variables:

- If the organism has regenerative potential and adult tissues are to be used, the temporal kinetics of the mitotic peak(s) following amputation may differ from planarians.
- Head/trunk fragments with mitotic cells and/or embryos can be used for planarian species with different regenerative capacities from those examined here.
- Puncturing the tissue with needles to break the external epithelium aids the penetration of DI H2_O_ into the tissues and cells. The duration of DI H2_O_ incubation, the size of the tissue fragment, the number of perforations made in the tissue, and the colchicine concentration and incubation time are all critical steps to optimize.
- The pressure applied to the coverslip during the squash is correlated to both size and texture of the tissue fragment. If the applied pressure is too great, the chromosomes may not remain grouped together. However, if the pressure is too minimal, nuclei or chromosomes may overlap.
- Imaging with a confocal microscope is not required, and a common compound epifluorescence microscope could be used as well.

### Immunohistochemical staining on planarians

Tail fragments were incubated in 5% N-Acetyl Cysteine (NAC) (Sigma Aldrich) in 1× PBS for 10 min at RT to remove mucous from the epithelium. The tissues were then fixed in pre-chilled Carnoy’s fixative solution (6:3:1 dilution of methanol:chloroform:glacial acetic acid) for 2 h at 4 °C. The samples were then rinsed in 100% methanol. The rehydrated tissues were bleached in 3% formamide and 6% H_2_O_2_ in 0.5% Triton X-100 in 1× PBS for 1 h under direct light. The blocking step in 5% horse serum in 0.3% Triton X-100 in 1× PBS for 1–2 h at RT was followed by the incubation ON in 1:500 Rabbit anti-phospho-Histone H3 (Ser10) Ab (H3P) (ab-32107, Abcam) at RT. The samples were rinsed and then incubated in 1:500 Goat anti-Rabbit IgG (H+L) Ab conjugated with Alexa Fluor^®^ 488 (ab-150081, Abcam) ON at RT. The samples were rinsed and mounted with ScaleA2 (4 M Urea, 0.1% Triton X-100, 20% Glycerol, 2.5% DABCO in ddH_2_O) mounting media.

Z-stack images were acquired with a Nikon Eclipse Ti microscope equipped with a Yokogawa W1 spinning disk head and robotic plate loader. Slides were loaded automatically and imaged with a low magnification objective. Objects were identified using custom software and imaged again with a Plan Apo 20× 0.75 NA air objective. Images were batch stitched and processed using custom plugins and macros in FiJi similar to previous reports [24].

### FISH protocol

Chromosome spreads were prepared according to the species-specific protocols described previously through the coverslip removal step. Once the slides were warmed to RT, they were rinsed 3 times with 1× PBS for 5 min each. The slides were then dehydrated in an ice-cold ethanol series (70%, 80%, and 100% ethanol) for 2 min each. Slides were then stored at RT for at least 2 days and as long as 1 month.

The telomere DNA probe (sequence: [TTAGGG]×7) was obtained from Integrated DNA Technologies and labeled with DIG-dUTP using the recombinant Terminal Transferase (Roche) according to the manufacturer’s protocol. For each slide, 20 ng of labeled telomere DNA probe was mixed with 10 volumes of Master Hybridization Mix (4× Saline-Sodium citrate buffer [SSC], 20% dextran sulfate, 2 mg/mL nuclease-free Bovine Serum Albumin [BSA], 50% deionized formamide in ddH2_O_). The total volume (~22 μl) was placed on the area containing the chromosome spreads, covered with a 22 × 22 mm coverslip, and sealed with mineral oil. The slides were incubated for 5 min at 70 °C to denature the DNA, followed by 24–36 h at RT for hybridization. Next, the slides were submerged in 2× SSC in a slide staining jar to remove the coverslips and then rinsed successively with 2× SSC, 0.5× SSC, and TNT (100 mM Tris-HCl, 150 mM NaCl, 0.1% Tween 20) for 15 min each at RT. The areas containing the chromosome spreads were covered with 1:200 anti-DIG-Rhodamine (Sigma Aldrich) and 1:1000 DAPI in TNB (5% Fetal Bovine Serum [FBS] in TNT) in the dark for either 1–4 h at RT or ON at 4 °C. The slides were rinsed with TNT for 15 min at RT and then mounted, imaged, and stored at 4 °C.

### Statistics analysis

An unpaired two-sample *t*-test was performed for calculation of the statistical difference between the number of H3P-positive cells in tail fragments at 0 hpa incubated with (N= 20) and without colchicine (N= 19) and at 6 hpa incubated with (N= 18) and without colchicine (N= 17). The differences were considered statistically significant with *P* < 0.05.

## Results

### Optimization of a chromosome preparation protocol

A simple protocol to study chromosome karyotypes has been optimized in three different organisms: the freshwater planarian *S. mediterranea*, the apple snail *P. canaliculata*, and the sea anemone *N. vectensis* (**Figure 1**). Briefly, adult tissues containing mitotic cells or entire embryos were collected and subjected to colchicine treatment. DI H_2_O was used to swell the cells of the tissues, which were then fixed in Carnoy’s fixative solution and moved onto a slide. Tissues were then treated with 60% acetic acid and squashed under a coverslip by manually applying pressure using the thumb. After an ON incubation at 4 °C, the coverslip was removed after snap-freezing with dry ice or liquid nitrogen. The chromosomes were then rinsed and stained with a DNA dye, or stored at RT for other experimental purposes.

**Figure 1.**
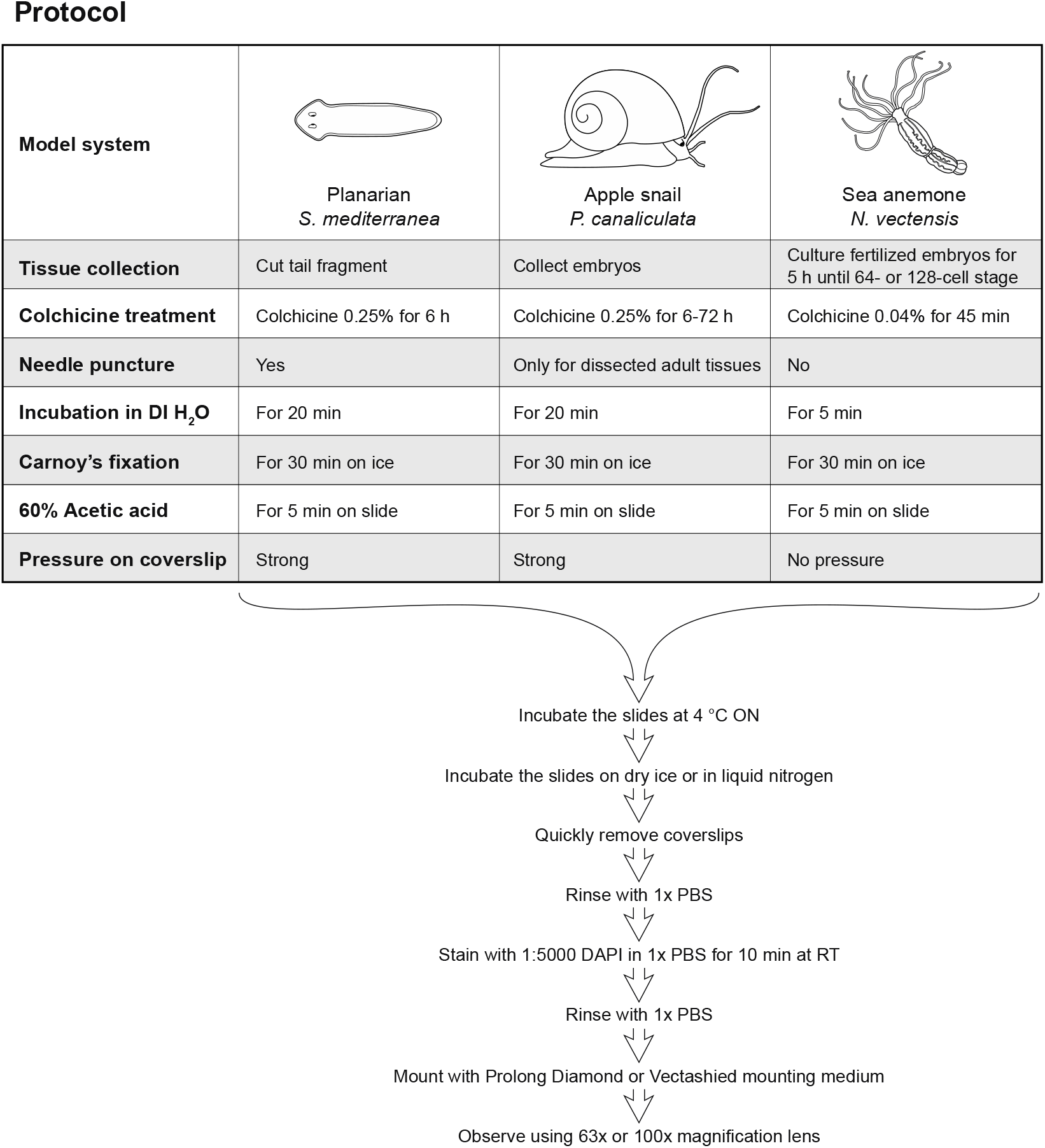
Schematic illustrating of the chromosome preparation protocols optimized for each model organism. Key steps in the chromosomal spread protocol are summarized as a flowchart for three invertebrate species. The first part of the protocol consists of steps that require species- and tissue-specific optimization. Each column highlights similarities and differences among the procedures for the planarian *S. mediterranea*, the freshwater snail *P. canaliculata* and the sea anemone *N. vectensis*. The second part of the protocol did not require species-specific optimization and was standardized for all organisms tested.

For freshwater planarians, tail fragments were used for karyotyping (**Figure 2**). In our experience, puncturing tail fragments with a needle while immersed in DI H_2_O was essential to obtain optimal and robust results. Punctures likely facilitate the penetration of DI H2_O_ into tissues, and practice was needed to achieve the best results. With more than 10 punctures in a 0.5 × 1 mm sized tail fragment, the wounds did not heal properly, which led to excessive exposure of the internal tissues to DI H_2_O (**Figure 2A, D**). These tissues fell apart during the 60% acetic acid treatment and yielded only a few intact chromosome spreads because the majority of chromosomes were scattered without discernable cell boundaries (**Figure 2I**). With few to no injuries, the tissues and cells of the tail fragment did not swell enough during DI H_2_O treatment (**Figure 2C, F**). Spreads from these tissues tended to have very crowded chromosomes (**Figure 2J**) or chromosomes that overlapped with interphase nuclei, making an accurate analysis of chromosome number and morphologies difficult. A properly punctured tail fragment increased slightly in size after DI H_2_O treatment (**Figure 2B, E**). It expanded more, but retained integrity after 60% acetic acid treatment (**Figure 2G**). A tail fragment could be sliced into several smaller pieces (250 × 250 μm) (**Figure 2H**), and this was sufficient to produce enough chromosome spreads with optimal morphologies. If the tissue used for the squash was too big, the chromosome spreads produced could be crowded and overlapping (**Figure 2J**). With optimal number of punctures, tissue swelling, tissue size and pressure applied on the coverslip, an optimal karyotype can be obtained (**Figure 2K, L**), where the chromosomes did not overlap or mix with chromosomes from other nuclei.

**Figure 2.**
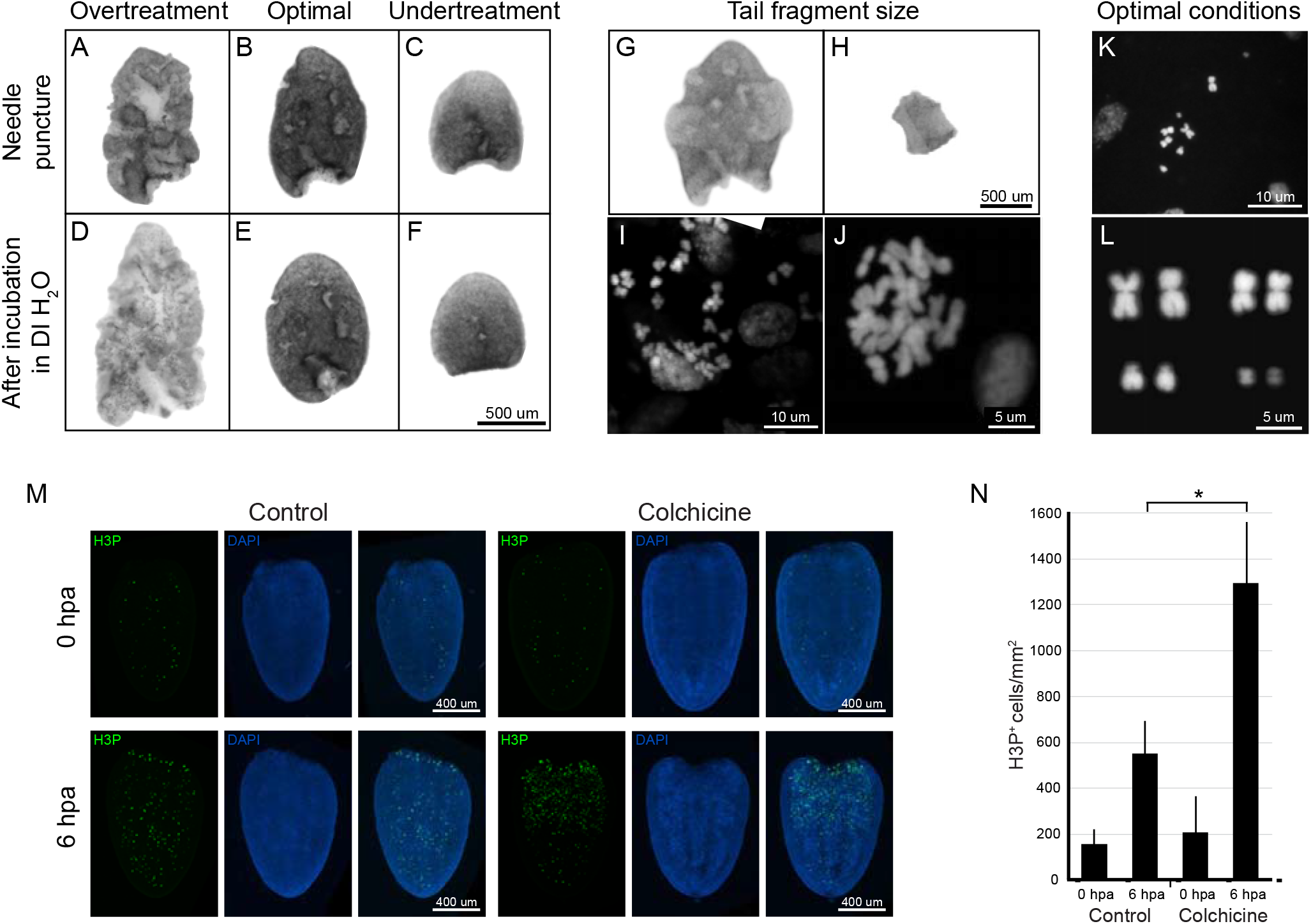
Protocol optimization in the freshwater planarian *S. mediterranea*. Conditions critical for optimal chromosome preparation in planarians include extent of needle puncture and colchicine treatment. An excessive number of needle punctures can cause deformation and disaggregation of the tissue, which became too swelled and fragile after incubation in DI H_2_O (**A, D**). This led to dispersed chromosomes on the slide without distinguishable cell boundaries (**I**). An optimal number of needle punctures (4~10 in a 0.5 × 1 mm fragment) facilitated tissue swelling without causing disaggregation in DI water (**B, E**) and maintained tissue integrity after acetic acid treatment (**G**). This produced optimal spreading of chromosomes on a slide (**K, L**). An insufficient number of needle punctures did not swell the tissues in DI H2_O_ (**C, F**), which led to overlapping chromosomes (**J**). One fourth of a 0.5 × 1 mm fragment (**G, H**) can produce sufficient optimal chromosome spreads. Too large a fragment used for squashing will also produce crowded chromosomes (**J**). Colchicine treatment increased the number of mitotic cells by ~2.5 times (**M, N**, \**P* < 0.0001). Mitotic cells were labeled by anti-H3P Ab (**M**).

Treatment with the mitosis-inhibiting drug colchicine significantly increased the number of mitotic cells in planarians (**Figure 2M, N**), similarly to other systems [25]. When colchicine binds tubulin, it prevents spindle formation and halts dividing cells in metaphase. An antibody detecting phospho-Histone H3 (Ser10) (H3P) was used to assess the number of mitotic cells in a planarian tail with or without 6 h of colchicine treatment. In the absence of colchicine, the number of dividing cells approximately tripled at 6 h post amputation (hpa) compared to baseline (0 hpa). Hence, with colchicine treatment, the number of mitotic cells increased about 7 times by 6 hpa compared to baseline, yielding a significantly higher number of chromosome spreads in the tissue (**Figure 2N**).

### Applicability of the protocol to multiple planarian species

To survey genome heterozygosity and diversity in populations of *S. mediterranea*, wild planarians were collected in the field in Sardinia (Italy) [19] (**Figure 3A**). Under a dissecting microscope, we defined three different morphological groups based on the shapes of the head and eyes, and the body pigmentation (**Figure 3B**). These features, however, were not sufficient to discern distinct species. We used the chromosome preparation protocol optimized on *S. mediterranea* to karyotype the planarian species present in this wild-caught population. Planarians in the second morphological group were diploid with 2n=8 chromosomes (**Figure 3C, E**), while those in the first morphological group were triploid with 3n=12 chromosomes (**Figure 3D, E**). Hence, we were able to readily determine that animals belonging to the second morphological group were *S. mediterranea* because they had the same karyotype as previously documented (**Figure 2K, L**) [26].

**Figure 3.**
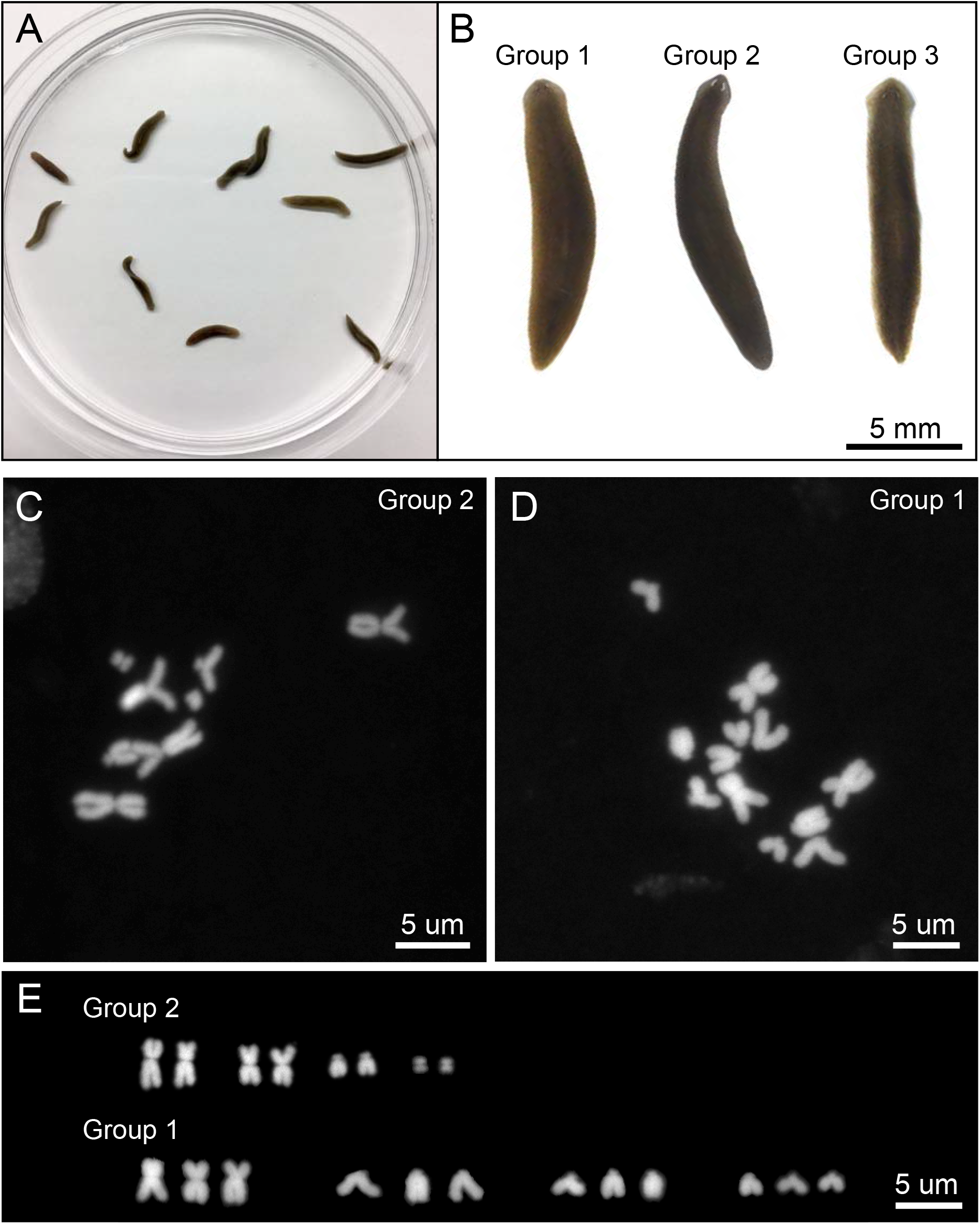
Species identification through karyotyping of wild planarians. Clearly distinct karyograms were obtained from three groups of morphologically similar planarians obtained in the wild. **A**: A mixed population of planarians collected in Sardinia (Italy). **B**: Wild planarians grouped based upon their gross morphology. **C**: Karyotype belonging to planarians in the second morphological group. This karyotype is the same as the *S. mediterranea* karyotype. **D**: Karyotype for planarians belonging to the first morphological group, which is *Schmidtea polychroa*. **E**: Karyogram of the planarians belonging to the second and first morphological group, which are diploid and triploid organisms, respectively.

This optimized and simple karyotyping protocol also works in planarian species other than *S. mediterranea*. First, we applied the protocol to *Dugesia japonica*, a planarian species reported to have mixed ploidy in different cells of the same animal [27] (**Figure 4A**). Indeed, during our analysis, multiple individuals were identified with mitotic cells that were either diploid (2n=16) (**Figure 4B**) or triploid (3n=24) (**Figure 4C**). The co-existence of diploid and triploid adult dividing cells in the same animal was intriguing. We quantified the ratio of diploid to triploid mitotic cells in *D. japonica*. Interestingly, there were consistently more diploid cells than triploid cells in all four animals examined, and the ratio varied from 2.30 to 3.46 with an average of 2.8 (**Figure 4D**).

**Figure 4.**
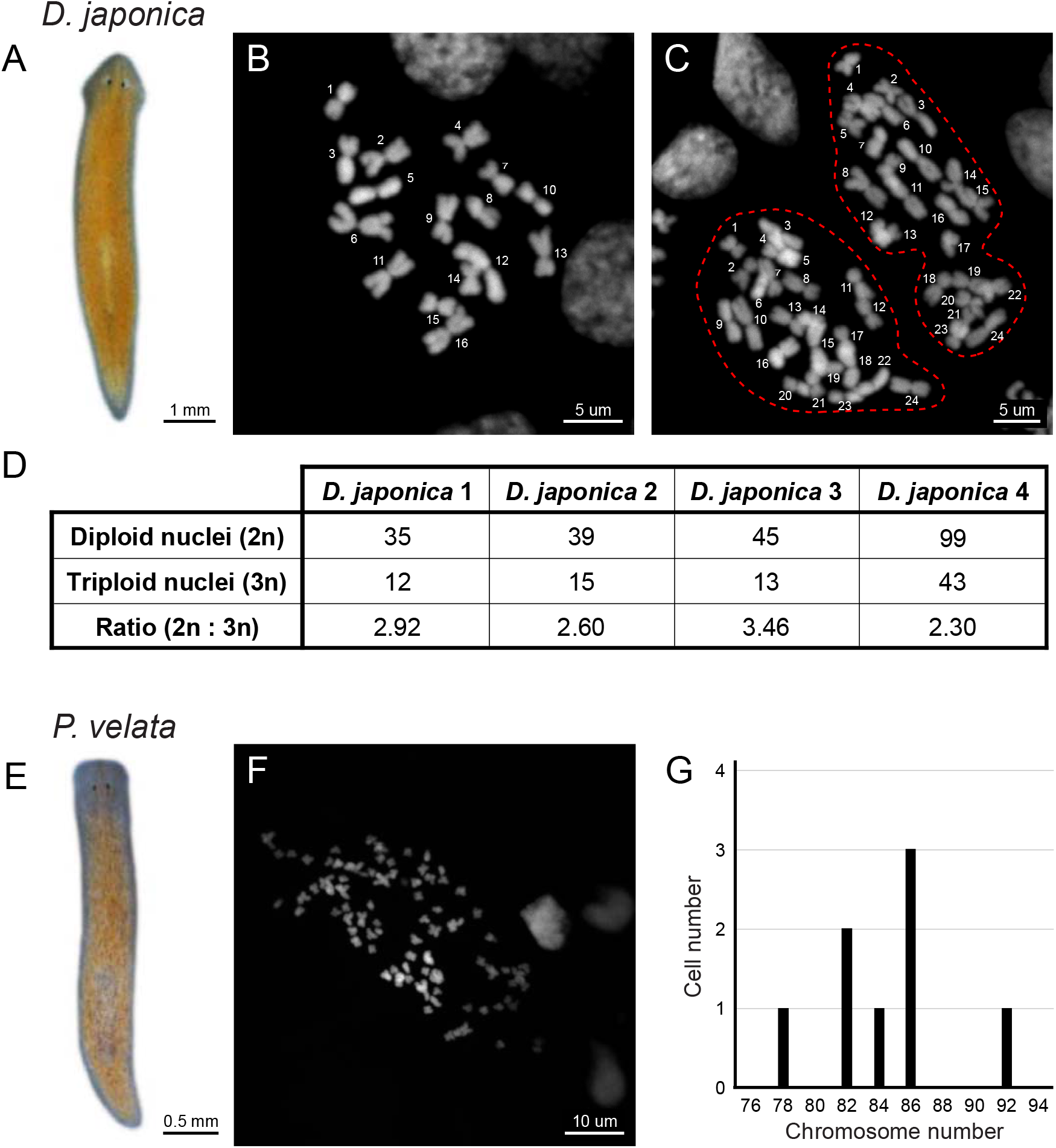
The optimized chromosome preparation protocol allows karyotyping of multiple planarian species. The optimized chromosome preparation protocol was used to generate karyotypes for two additional planarian species. **A**: *Dugesia japonica*. **B, C**: Karyotypes obtained for *D. japonica* demonstrating the presence of both diploid (**B**) and triploid (**C**) cells in the same individual. **D**: Table summarizing the ratio of diploid to triploid dividing cells in four *D. japonica* individuals. **E**: *Phagocata velata*. **F, G**: A representative *P. velata* karyotype (**F**) and quantification of the number of chromosomes per cell (**G**).

Second, we applied the protocol to *Phagocata velata*, a fissiparous species, that reproduces asexually by fission [28] (**Figure 4E**). The protocol worked successfully on its tail fragments, and we observed that this species has the largest number of chromosomes in planarians reported thus far (**Figure 4F**). The specific number of chromosomes seems to vary from cell to cell, but, on average, there are about 84 chromosomes per nucleus (**Figure 4G**). Altogether, the method reported here was successful in rapidly generating unambiguous karyotypes for a diverse cohort of planarian species.

### Applicability of the protocol to species belonging to different phyla

The planarian optimized protocol was successfully applied to the apple snail *P. canaliculata*, with minimal modifications. To obtain chromosomes for karyotyping, we first applied this method on embryos 5 days post fertilization (5 dpf), which have tissues undergoing abundant and frequent cell divisions. We found that, in the absence of colchicine treatment, dividing cells in various phases of mitosis were visible, but chromosomal compaction was not uniform (**Figure 5A–C**). Optimal chromosome spreads are usually obtained from cells in metaphase with highly condensed chromosomes. Therefore, we concluded that for karyotyping *P. canaliculata*, colchicine treatment was also required. As in planarians, a treatment with colchicine for 6 h in 5 dpf embryos overcame the problem of insufficient chromosome compaction and yielded many cells blocked in metaphase that had well-condensed chromosomes (**Figure 5D**). The analysis of these spreads confirmed reports by Mercado Laczkó and Lopretto [29] and Von Brand et al. [30], who each performed karyotype analyses on this species. As shown previously, we found that *P. canaliculata* nuclei were diploid and the number of chromosomes was consistent (2n=28 chromosomes). Moreover, the karyogram confirmed the presence of a pair of sexual chromosomes [30] (**Figure 5E**).

**Figure 5.**
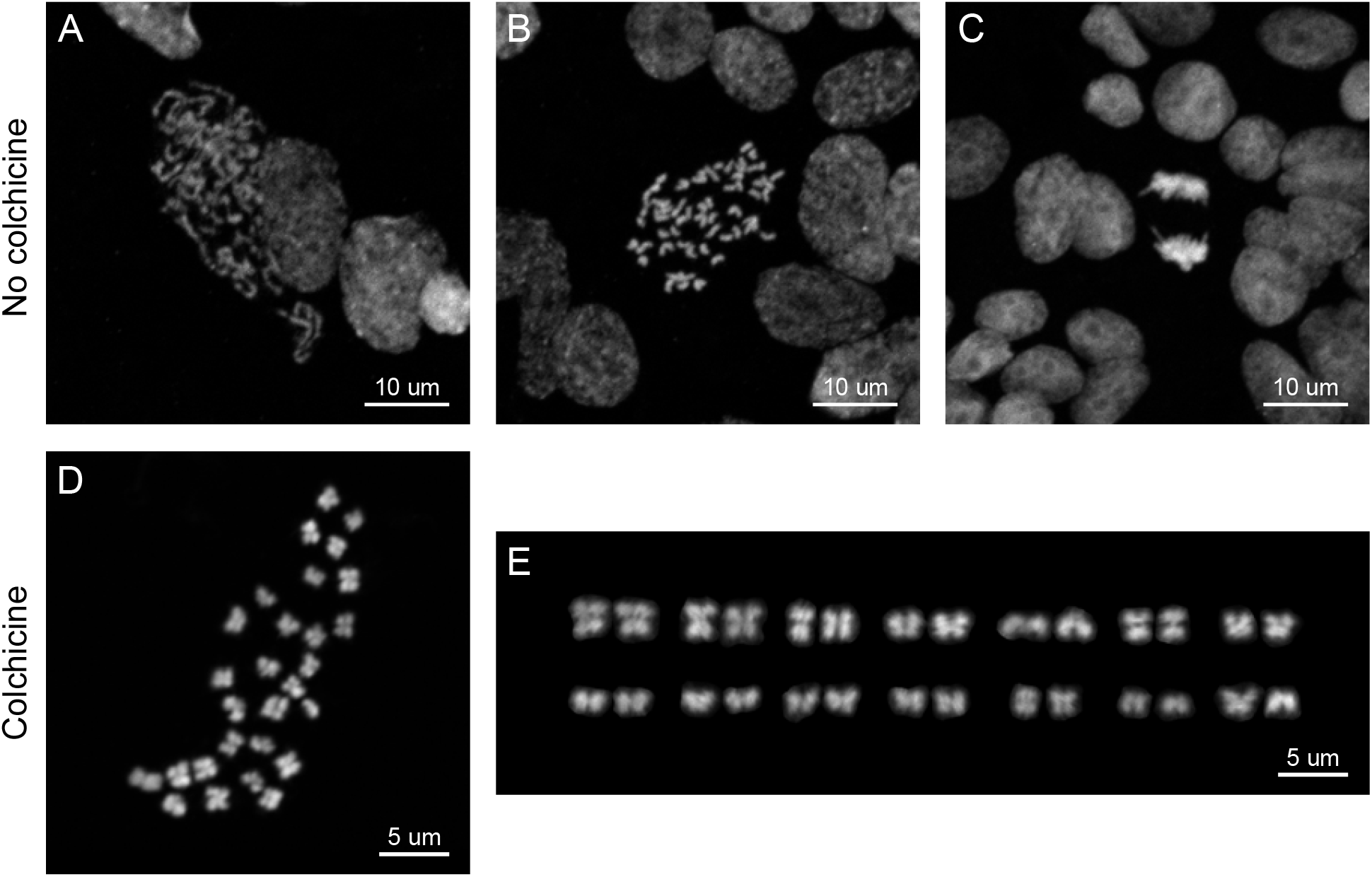
Chromosome preparation in the freshwater snail *P. canaliculata*. The optimized chromosome preparation protocol in planarians was used to generate karyotypes for both adult and embryonic tissues of the freshwater snail *P. canaliculata*. **A-C**: Mitotic cells showing (**A**) early prophase, (**B**) late prophase or metaphase, and (**C**) anaphase in embryos 5 dpf without colchicine treatment. **D, E**: (**D**) Karyotype and (**E**) karyogram obtained from *P. canaliculata* embryos 5 dpf treated with colchicine. They are also representative of chromosomes spreads obtained from the anterior and posterior kidney, and digestive gland in adult snails.

To further test the adaptability of this protocol for karyotyping using different tissues, we used it on adult organs of *P. canaliculata*. After an incubation for 72 h in colchicine the snails were dissected to collect organs known to undergo rapid cell turnover including gonads, gills, gut, anterior and posterior kidney, and digestive gland. We obtained good chromosome spreads from the anterior and posterior kidney, and the digestive gland (representative karyotype and karyogram are shown in **Figure 5D, E**). The tissue producing the most chromosome spreads were the gonads. However, if the gonads are to be used for karyotyping studies, it is crucial to be aware that this tissue contains many dividing polyploid cells (unpublished data). Shorter colchicine incubation (6, 24 and 48 h) provided none or extremely rare mitotic cells in the analyzed adult tissues.

Since the protocol successfully worked on both 5 dpf embryos and adult tissues of *Pomacea* by varying time of colchicine treatment, we tested how adaptable this protocol is for other embryonic stages. We asked whether a good karyotype can be obtained from any source of tissues by simply adjusting the time of the colchicine incubation (**Figure 6**). Embryos 5, 6, 8, and 10 dpf (**Figure 6A, D, G, J**) were treated with colchicine for 6 h and used for chromosome preparation. We observed that younger embryos provided more spreads, but the density of dividing cells was too high in certain areas of the slide, making it difficult to distinguish cell boundaries (**Figure 6B**). The older embryos, on the other hand, had lower density of mitotic cells (**Figure 6E, H, K**). We concluded that embryos 6 to 10 dpf treated with colchicine for 6 h produced enough chromosome spreads that were sufficiently separated from one another to easily associated them with individual cells. A longer colchicine treatment was also tested. After 24-h colchicine treatment, the chromosomes of 5 and 6 dpf embryos were found to be crowded and cell boundaries were hard to define. At these stages, 24-h colchicine treatment would be useful in cases where large numbers of chromosomes are needed, such as for chromosome sorting (**Figure 6C, F**). For older embryos (8 and 10 dpf), 24-h colchicine treatment turned out to be optimal for karyotype analysis (**Figure 6I, L**). Hence, a general rule of thumb in chromosome preparation in snails is that the older the tissue, the longer the colchicine treatment must be.

**Figure 6.**
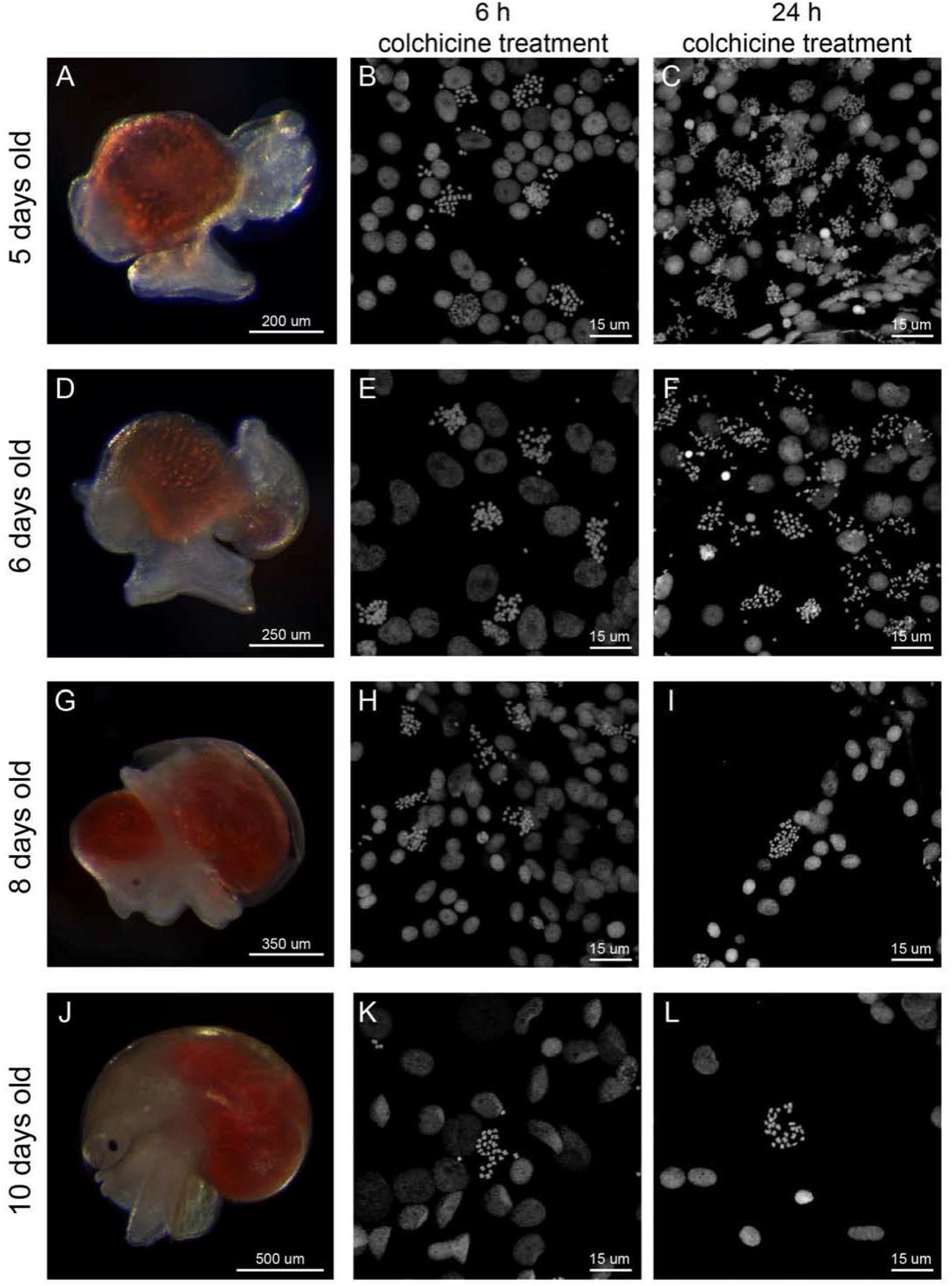
Chromosome spreads for different embryonic stages of *P. canaliculata*. The duration of colchicine treatment on embryos resulted in different densities of mitotic chromosomes for different embryonic stages. **A, D, G, J**: *P. canaliculata* embryos 5, 6, 8, and 10 dpf respectively, after removal of egg shell and perivitelline fluid. **B, E, H, K**: Chromosome spreads obtained from embryos at indicated developmental stages incubated in colchicine for 6 h. **C, F, I, L**: Chromosome spreads obtained from embryos at indicated developmental stages incubated in colchicine for 24 h.

Next, we tested the chromosome preparation protocol in a phylogenetically more distant species, the cnidarian *N. vectensis* (**Figure 7A**). Embryos between 64- and 128-cell stage were chosen as sources of mitotic cells (**Figure 7B, C**). At this stage, cell divisions are still synchronous with a 45-min interval. We showed that the best chromosome spreads were obtained with a 45-min treatment with 0.04% colchicine (**Figure 7D, E**). Nearly all blastomeres were arrested in metaphase and displayed well-condensed chromosomes. A 5-min osmolarity shock in DI H_2_O and a quick fixation using Carnoy’s fixative solution provided sufficient separation between individual chromosomes. Up to 10 fixed embryos were placed on one glass slide with a coverslip gently placed on top. The weight of the coverslip naturally flattened the embryos, achieving minimal overlapping between neighboring cells (**Figure 7H**). When the DI H_2_O treatment was too long or when additional pressure was applied to the coverslip, we observed frequent chromosome loss, as well as alterations in chromosome morphology in the final spread (**Figure 7F, G**). In contrast to other species examined in this study, *N. vectensis* sister chromatids tended to remain in close proximity to each other, making the chromosomes appear “bar-shaped” instead of a more classic “X” shape (**Figure 7D, E**). Subsequent quantification of chromosome numbers in 48 blastomeres confirmed the observation reported by Putnam et al. [31], who showed that *N. vectensis* nuclei are diploid (2n=30) (**Figure 7I**).

**Figure 7.**
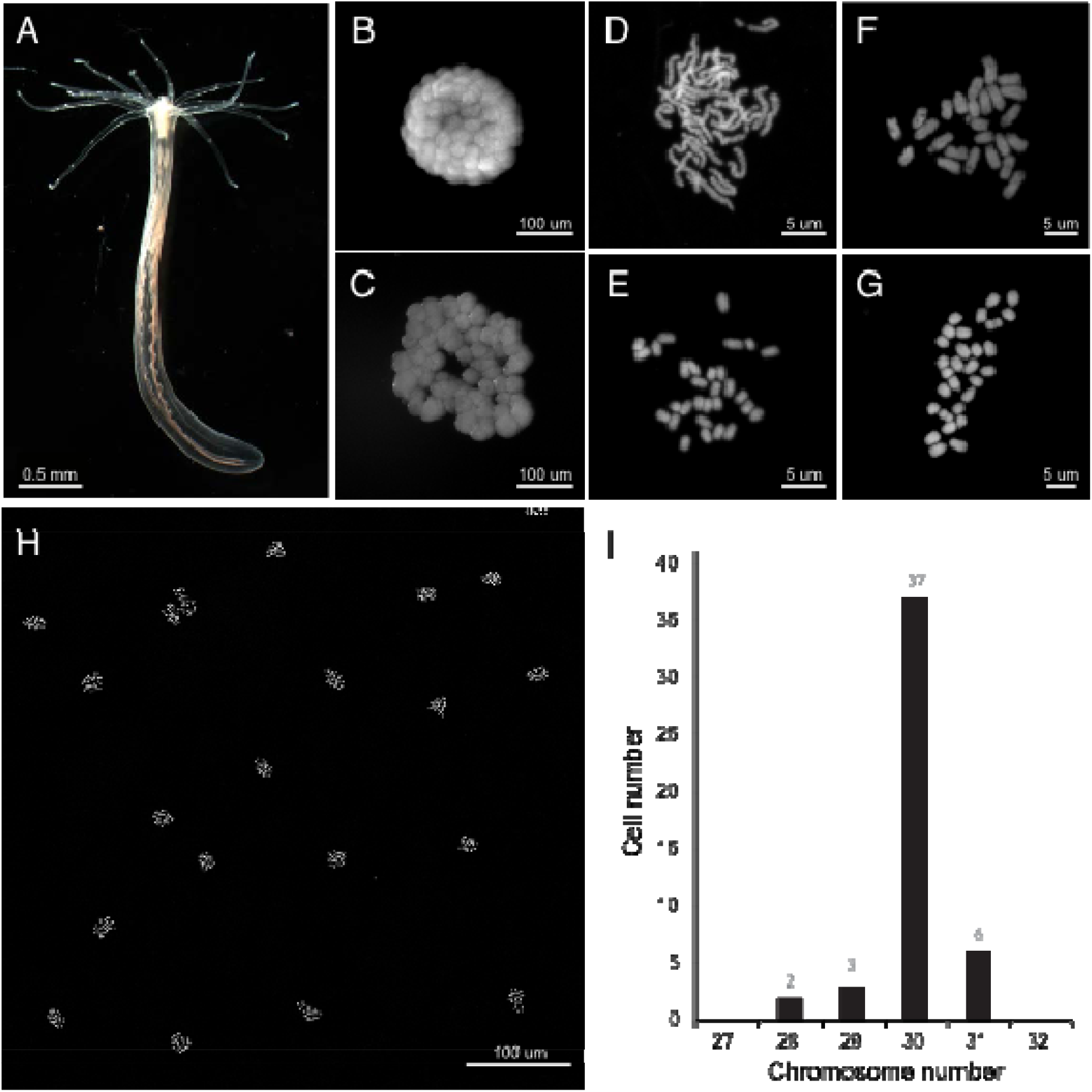
Chromosome preparation in the embryos of the sea anemone *N. vectensis*. The optimized chromosome preparation protocol was used to generate karyotypes for embryos of the sea anemone *N. vectensis*. **A**: Juvenile stage of *N. vectensis*. **B, C**: A 128-cell embryo (**B**) before and (**C**) after 5 min DI H_2_O. **D**: Karyotype obtained from *N. vectensis* embryos after a 30-min incubation in 0.04% colchicine. Notice the incomplete condensation of chromosomes. **E**: Karyotype obtained from *N. vectensis* embryos after a 45-min incubation in 0.04% colchicine. (2n=30). **F, G**: Excessive DI water incubation or physical force during preparation results in alteration of chromosome morphology and chromosome loss (**F**: 2n=25, **G**: 2n=29). **H**: Chromosome spreads obtained using 128-cell embryos imaged under lower magnifications. All blastomeres were synchronously arrested at metaphase. **I**: Quantification of the number of chromosomes per cell.

### Chromosome preparation protocol is compatible with downstream applications

Chromosome spread preparation can serve as the first step for other downstream protocols, such as chromosomal fluorescent *in situ* hybridization (FISH). Chromosomal FISH aims to localize the physical positions of genes and/or mutations on chromosomes.

To test whether our protocol can provide chromosomes suitable for further analyses other than the assembly of karyograms, we used highly conserved telomere-specific DNA probes labeled with DIG-dUTP (sequence: [TTAGGG]_n_) [32] for FISH. After the preparation of chromosomes following the protocol described in this paper, chromosomal FISH was performed (**Figure 8**). Specific and intense fluorescent signals at the ends of each sister chromatid were observed in all chromosomes obtained from adult *S. mediterranea* (**Figure 8A**), *P. canaliculata* embryos 6 dpf (**Figure 8B**), and 128-cell stage embryos of *N. vectensis* (**Figure 8C**).

**Figure 8.**
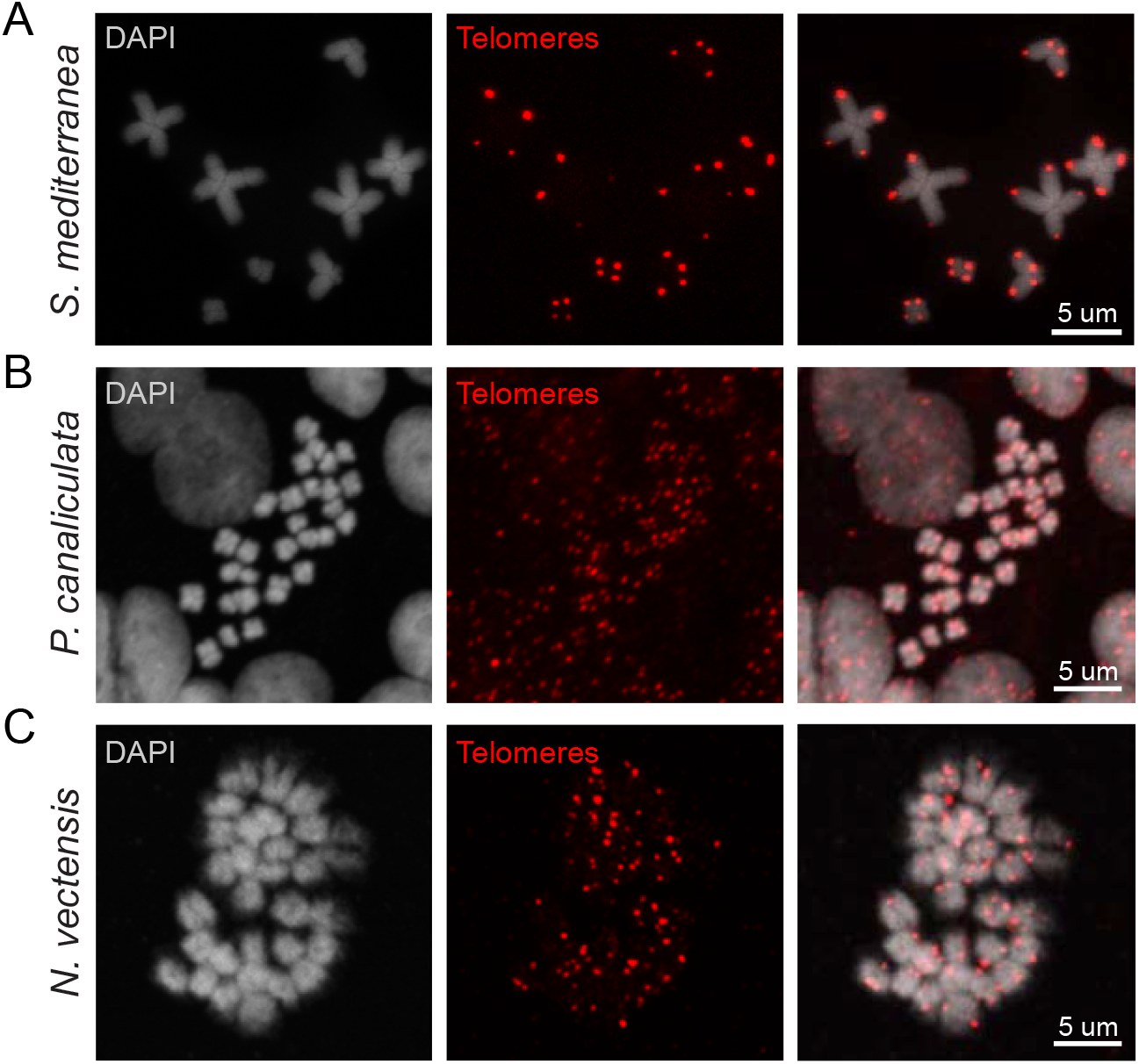
Telomere FISH on chromosome spreads of three model organisms. The chromosome preparation protocols optimized for each species are compatible with chromosomal FISH. Chromosomes spreads obtained from **A**: *S. mediterranea*, **B**: *P. canaliculata*, and **C**: *N. vectensis* were successfully stained with telomere-specific DNA probes labeled with DIG-dUTP.

## Discussion

Techniques providing easy and inexpensive access to chromosome composition (*i.e*., number and morphology) is invaluable for our understanding of chromosome behaviors and genetic architecture, particularly in understudied organisms. Here we reported an optimized protocol to prepare chromosomes for karyotyping and FISH in species from three different phyla: Platyhelminthes, Mollusca and Cnidaria. Several species from these related phyla are garnering the attention of communities interested in evolutionary developmental biology and evolutionary and population genetics. We expect that the protocol reported here will be highly adaptable to the study of multiple invertebrate species, and therefore be useful in studies of norms associated with chromosome composition, development, genetics, genomics and evolution.

The optimized protocol works efficiently in multiple planarian species, a freshwater apple snail and the starlet sea anemone. The reagents are highly accessible and inexpensive. Only a few steps of the protocol need to be adapted to specific features of the species of interest. Both adult tissues with dividing cells and embryonic tissues can be used as sources of mitotic chromosomes. Adjusting the treatment time with colchicine and DI water increases the rate of success, making this protocol highly adaptable to different tissues and different organisms. With this optimized protocol, we were able to confirm the karyotypes of all species examined. We were also able to visualize chromosome telomeres by FISH, demonstrating that chromosomes prepared with this protocol allow downstream applications.

In planarians, the analysis of karyotypes across multiple species was pioneered by Benazzi and his coworkers [18], and spanned from the early to late 20^th^ century. This body of work identified a large cohort of sexual and asexual planarian species and provided information on both the karyotypes and anatomy of sexual reproductive systems. However, few detailed karyotyping methodologies are available in planarian literature. Traditional methods employed regenerating blastemas 3 dpa for the preparation of chromosome spreads [33], which provided very little material to work with as the blastema are usually rather small. Furthermore, the traditional methods required a long processing time. Another issue associated with this literature is that the conditions for chromosome preparation are highly variable from researcher to researcher. Here, we showed that a fragment of tissue amputated from adult worm, and treated with colchicine for 6 h and successively with DI water as hypotonic reagent is sufficient to yield well-spread chromosomes in multiple planarian species. This can potentially function as a standardized methodology for planarian chromosome preparation.

To test the adaptability of the chromosome preparation methodology optimized on planarians, we applied it also to the freshwater apple snail *P. canaliculata* (Gastropoda, Ampullariidae) and the starlet sea anemone *N. vectensis* (Cnidaria).

*P. canaliculata*, originally from South America, is listed among the 100 most invasive species worldwide (Global Invasive Species Database), and the interest in this organism is significantly rising among researchers for both characterizing its immune system to limit its diffusion [34–36] and establishing it as a molluscan research organism for laboratory research [37–39]. We confirmed that the *P. canaliculata* genome is diploid with 2n=28 chromosomes [29, 30] applying the protocol optimized on planarian on both embryos at different developmental stages and adult tissues as source of mitotic cells. It is important to highlight that only a small number of optimizations were required to successfully isolate a significant number of good-quality chromosome spreads.

*Nematostella* has gained attention over the last decade due to both its unique phylogenic position as an early branching metazoan, and its amenability to experimental manipulation. The sequencing of the *Nematostella* genome in 2007 provided crucial insights into the ancestral eumetazoan gene repertoire and evolutionarily conserved signaling pathways [31]. In contrast to its apparent morphological simplicity, *Nematostella* possesses a complex genome [40, 41], whose sequence and high-quality annotation have provided a robust platform for applying modern molecular biological tools such as genome editing to the study gene function in the context of evolution [42, 43]. Yet, despite the rapid expansion of *Nematostella* as a new research organism, little attention has been paid to its chromosomal structure and organization. Previous scattered experimental evidence showed that *Nematostella* is a diploid animal with chromosome number 2n=30 [31]. Interestingly, this number is identical to several other hydrozoans and is also comparable to several species of *Acropora* corals (2n=28) [44, 45]. Utilizing the karyotyping method discussed above on early stage embryos (64- and 128-cell stage) and optimizing the incubation time in colchicine and DI water, and the pressure applied on the coverslip, we easily obtained highly synchronized metaphase spreads. Our data confirmed previous observations regarding chromosome number and morphology for *Nematostella*.

Karyograms have been used as a powerful tool for species identification, both among planarians and mollusks, for example [21, 46]. We showed that when planarians are collected in the wild, it can be difficult to distinguish them only on the basis of morphological features. The high regenerative potential of these animals allows the researcher to karyotype all the individuals without sacrificing them. Historically, planarian species identification relies on the anatomy of sexual reproductive organs [18, 47, 48], which leaves the cohort of asexual planarian species largely uncatalogued. A proper characterization of the karyotypes of both sexual and asexual species would facilitate the identification and cataloging of planarian species, providing, at the same time, a rich resource for the understanding of karyotype evolution [21]. Similarly, over the years, the Gastropoda class has been re-organized multiple times in subclasses and families, and new species have been integrated, but the lack of a unique and established method for species identification has created multiple incongruences [49, 50]. In particular, Hayes and colleagues faced this problem within the Ampullariidae family, a group of large freshwater snails, suggesting a combination of morphological and molecular features for identifying snail species [51, 52]. The karyogram (chromosome number and size, and position of centromere and G-bands) can be a very useful feature for species identification [29, 46, 53, 54]. The possibility to easily obtain unambiguous karyotypes from material collected in the field irrespective of embryonic developmental stages, or from adult tissue without extensive optimization or requiring many animals stands to be extremely useful in comparative and population genetic studies.

In conclusion, we optimized a chromosome preparation protocol in invertebrate species from three phyla. The protocol is simple, inexpensive and highly adaptable to multiple species. Prepared chromosomes can be used for multiple downstream applications including FISH for localizing telomeres and potentially any other gene of interest.

## Declarations

### Ethics approval and consent to participate

Studies with planarians, snails and sea anemone in here presented work abide to ethical regulations.

### Consent for publication

Not Applicable.

### Availability of data and material

Original data underlying this manuscript can be accessed from the Stowers Original Data Repository at http://www.stowers.org/research/publications/libpb-1241.

### Authors’ contributions

LG established the chromosome preparation protocol on planarians, and was a major contributor in writing the manuscript. AA established the chromosome preparation protocol on apple snails, and was a major contributor in writing the manuscript. SH established the chromosome preparation protocol on *Nematostella*, and contributed in writing the manuscript. CGH collected part of the data on planarians. SS contributed in screening already published protocols for karyotyping and testing different variables on both planarians and *Nematostella*. SM acquired images after H3P Ab staining and quantified the number of H3P-positive cells. MG and ASA funded the research and helped write the manuscript. All authors read and approved the final manuscript.

## Acknowledgements

We thank S.A. Elliott and S. Nowotarski for helpful comments on the manuscript; P. Newmark and O. Guedelhoefer for earlier work on the planarian karyotyping protocol; M. Miller for illustration assistance; the Stowers Aquatics Facility for maintaining all animals used in this study. We also thank members of Sánchez Alvarado and Gibson Laboratories for helpful comments and discussions and Sebastian Sánchez-Piotrowski for his assistance obtaining wild cultures of planarians.

